# Obesity/Type II Diabetes Alters Macrophage Polarization Resulting in a Fibrotic Tendon Healing Response

**DOI:** 10.1101/131607

**Authors:** Jessica E Ackerman, Michael B Geary, Caitlin A Orner, Fatima Bawany, Alayna E Loiselle

**Affiliations:** Center for Musculoskeletal Research, Department of Orthopaedics and Rehabilitation, University of Rochester Medical Center, 601 Elmwood Avenue, Box 665, Rochester, NY 14642

## Abstract

Type II Diabetes (T2DM) dramatically impairs the tendon healing response, resulting in decreased collagen organization and mechanics relative to non-diabetic tendons. Despite this burden, there remains a paucity of information regarding the mechanisms that govern impaired healing of diabetic tendons.

Mice were placed on either a high fat diet (T2DM) or low fat diet (lean) and underwent flexor tendon transection and repair surgery. Healing was assessed via mechanical testing, histology and changes in gene expression associated with collagen synthesis, matrix remodeling, and macrophage polarization. Obese/diabetic tendons healed with increased scar formation and impaired mechanical properties. Consistent with this, prolonged and excess expression of extracellular matrix (ECM) components were observed in obese/T2DM tendons. Macrophages are involved in both inflammatory and matrix deposition processes during healing. Obese/T2DM tendons healed with increased expression of markers of pro-inflammatory M1 macrophages, and elevated and prolonged expression of M2 macrophages markers that are involved in ECM deposition. Here we demonstrate that tendons from obese/diabetic mice heal with increased scar formation and increased M2 polarization, identifying excess M2 macrophage activity and matrix synthesis as a potential mechanism of the fibrotic healing phenotype observed in T2DM tendons, and as such a potential target to improve tendon healing in T2DM.

## Introduction

The dramatic increase in type II diabetes mellitus (T2DM) as part of the obesity epidemic (1), is one of the most critical health challenges facing the U.S.; in 2012 more than 29 million people, or nearly 10% of the US population were diabetic, resulting in a health care burden of $254 billion (2). T2DM results in systemic inflammation and is characterized by metabolic dysfunction including elevated plasma glucose levels (hyperglycemia). Among a plethora of systemic complications and co-morbidities arising from T2DM, the impact on the musculoskeletal system is emerging as an important disease manifestation. T2DM dramatically alters tendon homeostasis; diabetic tendons are generally fibrotic and display increased disorganization of the extracellular matrix (ECM) (3), although there is some variability in the degree to which different tendons are affected by T2DM (4, 5).

In addition to altering tendon homeostasis, T2DM dramatically impairs the tendon healing response (5, 6). Following injury, diabetic tendons heal with decreased mechanical properties (4, 7-9), and decreased collagen fiber organization (8), relative to non-diabetic controls. These changes in the healing cascade result in tendons that are more likely to rupture after repair and, in the case of flexor tendons (FTs), more susceptible to impaired gliding function due to the formation of fibrous adhesions between the tendons and surrounding tissue. In the hand, adhesions between the FT and synovial sheath impair gliding of the tendon and restrict digit range of motion (ROM), which in turn can impair the function of the entire hand (9). Considering that regaining satisfactory digital function in non-diabetic patients is one of the most pressing challenges facing hand surgeons (10), with 30-40% of primary FT repairs resulting in significant adhesion formation and loss of digit ROM (11), it is imperative to understand the healing process in the context of both non-diabetic and diabetic healing. However, there remains a lack of information regarding the cellular/molecular mechanisms of impaired tendon healing in both diabetic and non-diabetic patients. The cellular environment during tendon healing is complex and heterogeneous, and changes with the healing process through overlapping phases of inflammation, proliferation/matrix deposition, and remodeling. Among the cells involved, macrophages represent a rather dynamic population that, depending on phenotype, can modulate and participate in all phases of healing. Pro-inflammatory, classically activated M1 macrophages are involved in cell apoptosis, catabolism of ECM, and the inflammatory response. In contrast, alternatively activated M2 macrophages promote ECM deposition and restoration of native tissue architecture (12). Given the elevated inflammatory profile that exists in T2DM at baseline (13), and the ability for macrophages to modulate both the inflammatory response and subsequent matrix deposition/ remodeling, they represent a potentially important cell population in the impaired healing phenotype that occurs in diabetic tendon.

Previous studies using rodent models of T2DM have clearly defined impairments in FT mechanics as a major impediment to satisfactory healing (14), however, changes in gliding function were not assessed and the mechanism of these changes are unknown. In the present study we utilized a murine model of diet induced obesity and T2DM to test the hypothesis that obese/diabetic tendons heal with an increase in scar formation due to excess extracellular matrix deposition, relative to lean controls. Furthermore, we assessed macrophage content and polarization as a potential mechanism of altered healing in obese/diabetic tendons.

## Materials and Methods

**Ethics Statement:** This study was carried out in strict accordance with the recommendations in the Guide for the Care and Use of Laboratory Animals of the National Institutes of Health. The protocol was approved by University Committee on Animal Resources at the University of Rochester Medical Center (protocol #2014-004).

### Mice and Diet Induced Obesity/Type 2 Diabetes Model

Male C57Bl/6J mice (Jackson Laboratories, Bar Harbor, ME) or MaFIA (Macrophage associated Fas-Induced Apoptosis, #5070Jackson Laboratories, Bar Harbor, ME) were started on either a low fat diet (LFD; 10% Kcal, Research Diets #12450D) or high fat diet (HFD; 60% Kcal, Research Diets #D12492) beginning at 4-weeks of age, and were maintained on their respective diets for the duration of the experiment. Male C57Bl/6J mice were used as this strain and sex are very susceptible to diet-induced obesity and T2DM (8, 9). MaFIA mice express Green Fluorescent Protein (GFP) under the control of the *Csf1r*-promoter resulting in GFP expression in macrophages and dendritic cells. Mice were maintained in pathogen free housing on 12hr/12hr light/ dark cycle with ad libitum access to food and water (except during fasting when food was removed).

### Fasting Blood Glucose Measurement and Glucose Tolerance Testing

Mice were fasted for 6hrs (15). Fasting blood glucose (BG) was assessed immediately prior to glucose administration and was also used as the 0hr time-point in the glucose tolerance test (GTT). Mice were administered 2g/kg glucose in PBS via intraperitoneal injection, and BG was measured using a glucometer with blood from the tail vein. Glucose tolerance was assessed at 12-weeks post-diet initiation (16 weeks of age), two days prior to FDL surgery.

### Murine Flexor Tendon Healing Model

16-week old male mice (12 weeks after HFD or LFD initiation) underwent surgical transection and repair of the flexor digitorum longus (FDL) tendon as previously described (16). Animals were sedated using ketamine (60 mg/kg) and xylazine (4 mg/kg), and post-operative pain was managed with a single subcutaneous injection of 0.05 mL extended-release buprenorphine (1.3 mg/mL). The proximal FDL tendon was released at the myotendinous junction along the tibia in order to protect the distal repair after surgery. The FDL tendon was then exposed in the plantar hind foot using a longitudinal incision, and the tendon was transected in the mid-foot region. The tendon was immediately repaired using 8-0 nylon sutures (Ethicon Inc., Summerville, NJ) in a modified Kessler pattern. After repair, the hind foot and proximal release site were closed with a single 5-0 nylon suture (Ethicon Inc., Summerville, NJ). After the procedure, mice were allowed free active motion and weight bearing in their cages, and were monitored daily by veterinary staff.

### Adhesion Testing and Range of Motion

Animals were sacrificed between post-repair days 10 and 28 for adhesion testing (n=6-10 per group per time-point) as previously described (17, 18). Briefly, the proximal end of the FDL tendon was secured between two pieces of tape, and the hindpaw at the tibia was placed into a custom apparatus. The proximal FDL was sequentially loaded from 0 g to 19 g with images captured after each incremental load. Images were used to measure the flexion angle of the metatarsophalangeal (MTP) joint relative to the unloaded position. Measurements were made by two blinded, independent observers using ImageJ software (http://rsb.info.nih.gov/ij/).

### Biomechanical Testing

Immediately following adhesion testing, the FDL tendon was released from the tarsal tunnel. The proximal FDL and distal phalanges were secured opposite each other in grips in the testing apparatus (Instron 8841 DynaMight™ axial servohydraulic testing system, Instron Corporation, Norwood, MA), and displaced at 30 mm/minute until the tendon failed. The maximum load at failure, stiffness, yield load, and energy to max force were calculated from the automatically logged force-displacement data generated during testing.

### Histology and Immunohistochemistry

The hind limbs were harvested between days 7-28 post-repair as previously described (19) (n= 4-6 per group per time point). Briefly, the samples were fixed for 48 h in 10% neutral buffered formalin (NBF) with the tibia at 90° relative to the foot. Following fixation, decalcification and sucrose incubation the samples were embedded in OCT (Tissue-Tek, Sakura Finetek U.S.A., Inc., Torrance, CA) for frozen sectioning. Eight-micron sagittal sections were stained with alcian blue/hematoxylin and Orange G (ABHOG). Tendon repairs from MaFIA mice were harvested at day 21 post-repair (n=4 per group), fixed and decalcified as above, however they were processed and embedded for paraffin sectioning. Three-micron serial sections were stained with ABHOG, or underwent immunofluorescent (IF) or immunohistochemical (IHC) staining. Sections were probed with either Goat-anti-GFP (1:2500, #ab6693, Abcam, Cambridge MA), Rabbit-anti-F4/80 (1:500, #sc-26643, Santa Cruz Biotechnology, Dallas TX), Rabbit-anti-TNFα (1:100, #ab6672, Abcam, Cambridge MA), or Rabbit-anti-IL-1ra (1:10000, #ab124962, Abcam, Cambridge MA). For GFP IHC, sections were probed with biotinylated goat anti-rabbit secondary (#PK-4001, Vector Laboratories, Burlingame, CA) according to manufacturer instruction. Staining was visualized with DAB chromogen (#sk-4105, Vector Laboratories, Burlingame, CA), and counterstained with hematoxylin. For IF staining, sections were probed with donkey-anti-rabbit 594 fluorescently conjugated secondary antibody (#705-546-147, Jackson Immuno, West Grove, PA). DAPI was used as a nuclear counterstain.

### RNA Extraction and Real-Time RT-PCR

Immediately following sacrifice between 3-28 days post-repair (n=3 per group per time-point), sections of the FDL tendon containing the repair were carefully dissected. Tendons were pooled and homogenized using the Bullet Blender Homogenizer (#BB24-AU, Next Advance, Averill Park, NY) and Trizol reagent (Invitrogen, Carlsbad, CA). RNA (500ng) was reverse-transcribed using iScript (BioRad, Hercules, CA), and used as a template for real-time PCR with PerfeCTa SYBR Green SuperMix (Quanta Biosciences, Gaithersburg, MD) and gene specific primers (Table 1). Gene expression was standardized to the internal control *β*-*actin*, and normalized to expression at day 3 in LFD repairs. Data are presented as fold changes relative to LFD day 3 samples ± standard error of the mean (SEM). Samples were run in triplicates, and each experiment was repeated at least twice.

**Table 1.**
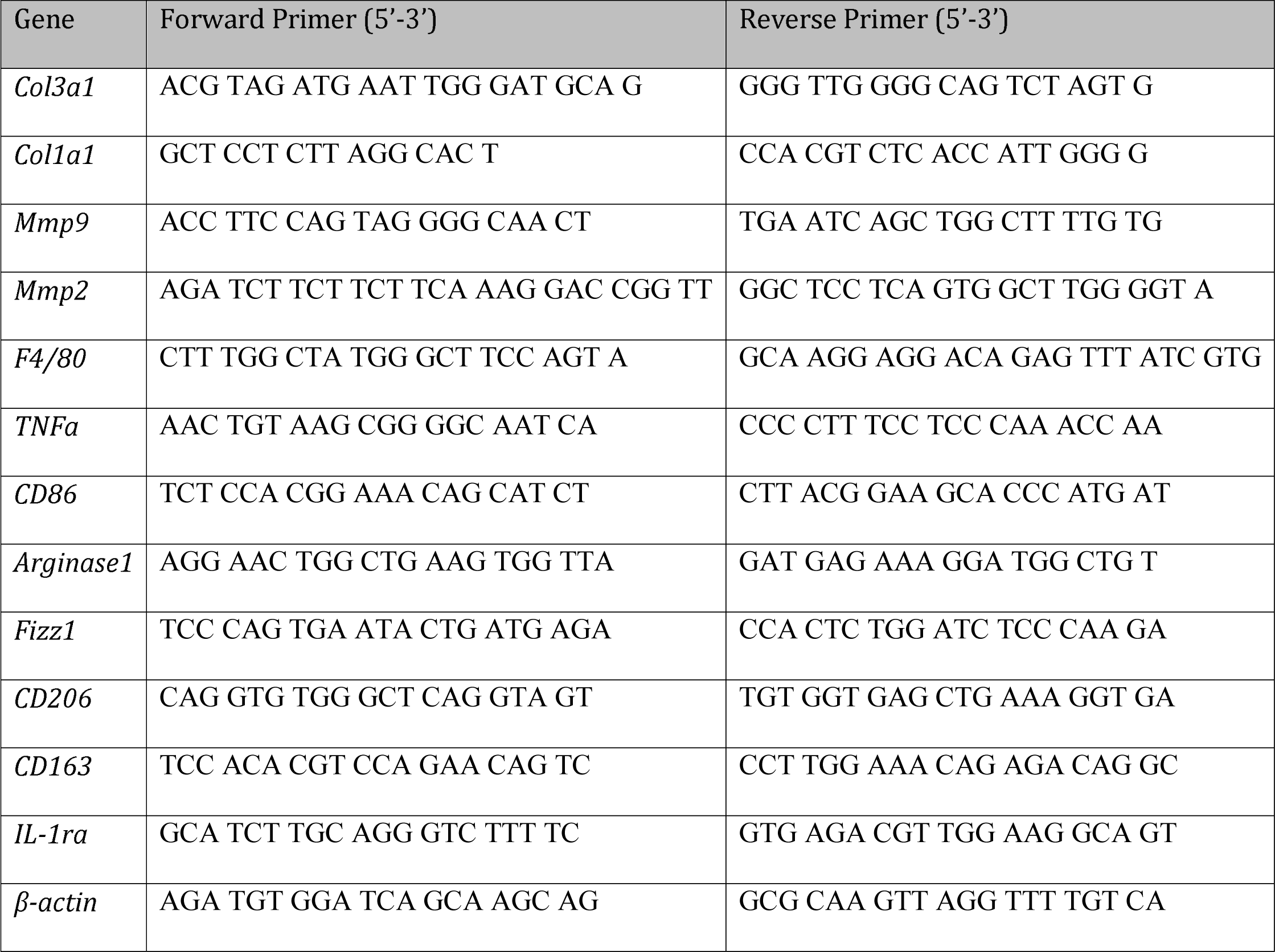
Primer sequences for quantitative PCR experiments.

### Statistical Analysis

Data are presented as mean ± SEM. Sample sizes were based on post-hoc power analyses of previous studies (17). Body weight and fasting BG levels were compared using one-way analysis of variance (ANOVA) with Bonferroni’s multiple comparisons test, with a single pooled variance. A two-way ANOVA with Bonferroni’s multiple comparisons test was used to analyze GTT, qPCR, biomechanical, and gliding data.

## Results

### HFD resulted in increased body weight and impaired glucose tolerance

Body weights of mice in the HFD and LFD groups were compared at the time of surgery (16 weeks of age, 12-weeks post-diet initiation) and the time of sacrifice. The HFD group had significantly higher body weight at both the time of surgery (LFD: 33.2g ± 0.6; HFD: 41.5g ± 1.3, p<0.05), and the time of sacrifice (LFD: 32.6g ± 0.6; HFD: 39.6g ± 1.2, p<0.05) (Figure 1A).

**Figure 1.**
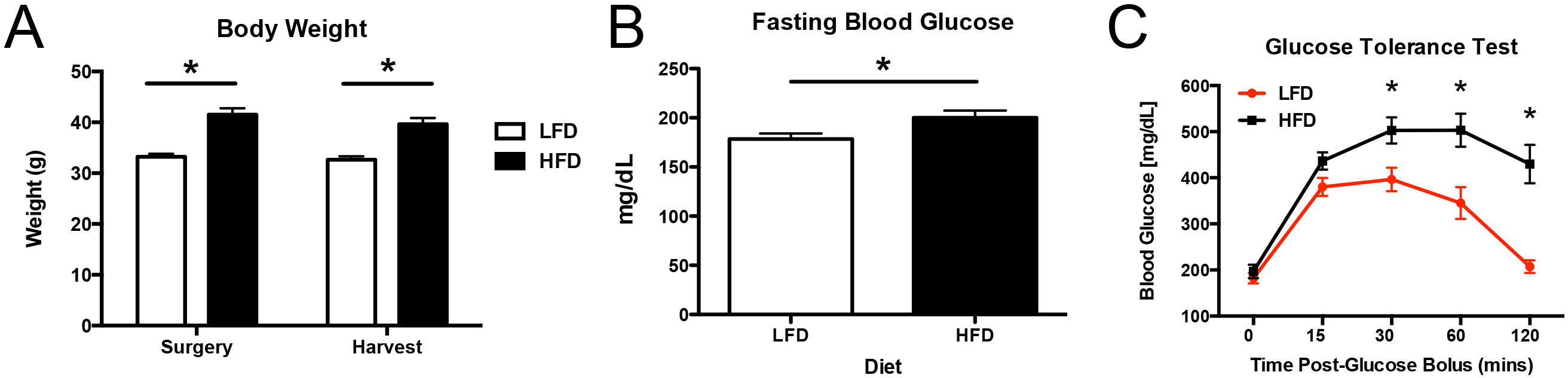
HFD leads to increased body weight and impairs glucose tolerance. A – Body weights from mice in both HFD (black bars) and LFD groups (white bars) were compared at the time of surgery and the time of harvest. At both points, the HFD group weighed significantly more than the LFD group. B – Elevated fasting blood glucose levels were observed in HFD (black bars), relative to LFD mice (white bars). C – Mice from the HFD group (black line) demonstrated an impaired ability to respond to glucose loading relative to LFD (red line), as shown by significantly elevated blood glucose levels at 30, 60, and 120 minutes after delivering a glucose bolus. (*) Indicates p<0.05.

Fasting BG was measured in both groups prior to surgery to confirm metabolic dysfunction and T2DM in HFD mice. BG was significantly increased in HFD mice, relative to LFD (LFD: 178.4 mg/dl ± 5.7; HFD: 200.2 mg/dl ± 7.3, p<0.05) (Figure 1B).

The ability to respond to glucose loading was assessed in both groups by GTT, with BG levels assessed up to 120 minutes after a standard glucose load. BG levels were significantly higher in the HFD group relative to LFD at 30 minutes (LFD: 396 mg/dl ± 25.8; HFD: 502.5 mg/dl ± 28.8, p<0.05), 60 minutes (LFD: 345.3 mg/dl ± 34.3; HFD: 503.3 mg/dl ± 36.3, p<0.05), and 120 minutes post-bolus (LFD: 207.3 mg/dl ± 13.8; HFD: 429.8 mg/dl ± 47.3, p<0.05) (Figure 1C).

### HFD impairs return to normal MTP flexion angle and gliding resistance

The MTP flexion angle offers a measure of the maximal flexion achieved at the highest load, with larger angles corresponding to greater flexion. The flexion angle showed substantial improvement at 28 days in the LFD group (129% increase over day 10, p<0.001) (Figure 2A). In contrast the MTP flexion angle in HFD repairs was not significantly improved relative to HFD day 10 repairs (31% increase, p>0.05). Moreover the flexion angle in the HFD group was significantly lower than the LFD group at 28 days (LFD: 44.6°± 3.9; HFD: 31.7°± 2.3, p<0.01) (Figure 2A).

**Figure 2.**
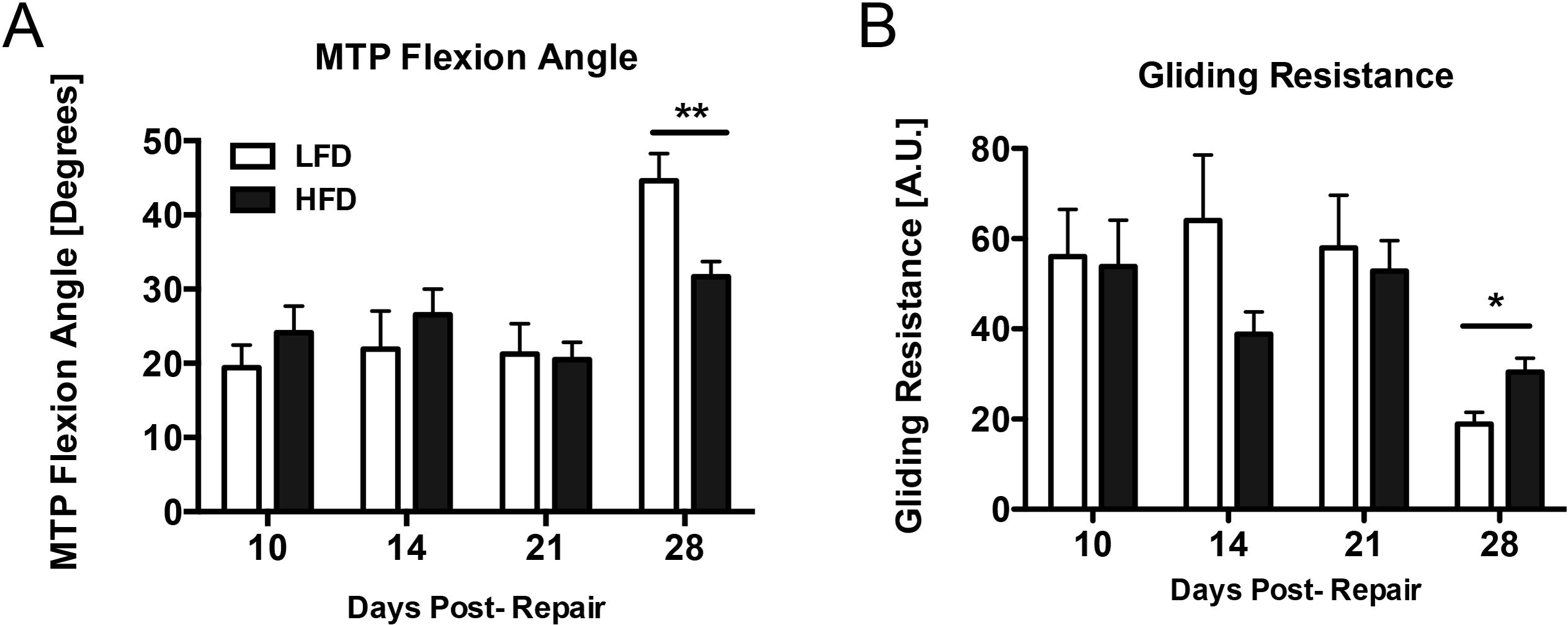
HFD impairs return to normal MTP flexion angle and gliding resistance. Incremental loading of the proximal end of the FDL was performed to measure the gliding resistance and the metatarsophalangeal (MTP) flexion angle in the hindpaw. A –LFD-fed mice (white bars) show substantial improvement in flexion angle 28 days post-repair, while the MTP flexion angle is significantly lower in the HFD group (black bars) at the latest time-point. B – Gliding resistance is significantly higher in the HFD group (black bars) 28 days post-repair relative to LFD (white bars), suggesting that T2DM interferes with the restoration of normal gliding and flexion after repair. (*) Indicates p<0.05.

Gliding resistance was determined by measuring the changes in MTP flexion with incremental loading of the tendon, with higher Gliding Resistance associated with increased scar formation. A decrease in Gliding Resistance was observed in both the LFD and HFD repairs between 10-28 days; however, Gliding Resistance was significantly increased in HFD repairs, relative to LFD at 28 days (LFD: 18.9 ± 2.5; HFD: 30.4 ± 3.4, p<0.05) (Figure 2B).

### Biomechanical properties are comprised in Obese/T2DM tendon repairs

Biomechanical testing was performed in both groups to determine the effects of T2DM on the mechanical properties of healing tendons over time. The maximum load at failure increased from 10 to 28 days in both the LFD (227% increase, p<0.0001) and HFD (125% increase, p<0.0001) (Figure 3A). At 28 days maximum load at failure was significantly lower in the HFD group relative to repairs in the LFD group (LFD: 2.9 N ± 0.1; HFD: 2.2 N ± 0.2, p<0.01) (Figure 3A). Tendon stiffness increased from 10 to 28 days in both LFD (215% increase, p<0.001) and HFD (223% increase, p<0.01) (Figure 3B), however no differences in stiffness were observed between HFD and LFD repairs at any time. Repairs from the HFD group had significantly lower yield load at 28 days relative to LFD repairs (LFD: 2.7 N ± 0.1; HFD: 1.2 N ± 0.1, p<0.01) (Figure 3C). The energy to maximum force increased at each time-point from 10 to 28 days in the LFD group, while no change occurred in HFD repairs between 21 and 28 days. Energy to maximum force was significantly lower at 28 days in the HFD group relative LFD (LFD: 0.9 N*mm ± 0.1; HFD: 0.5 N*mm ± 0.1, p<0.05) (Figure 3D).

**Figure 3.**
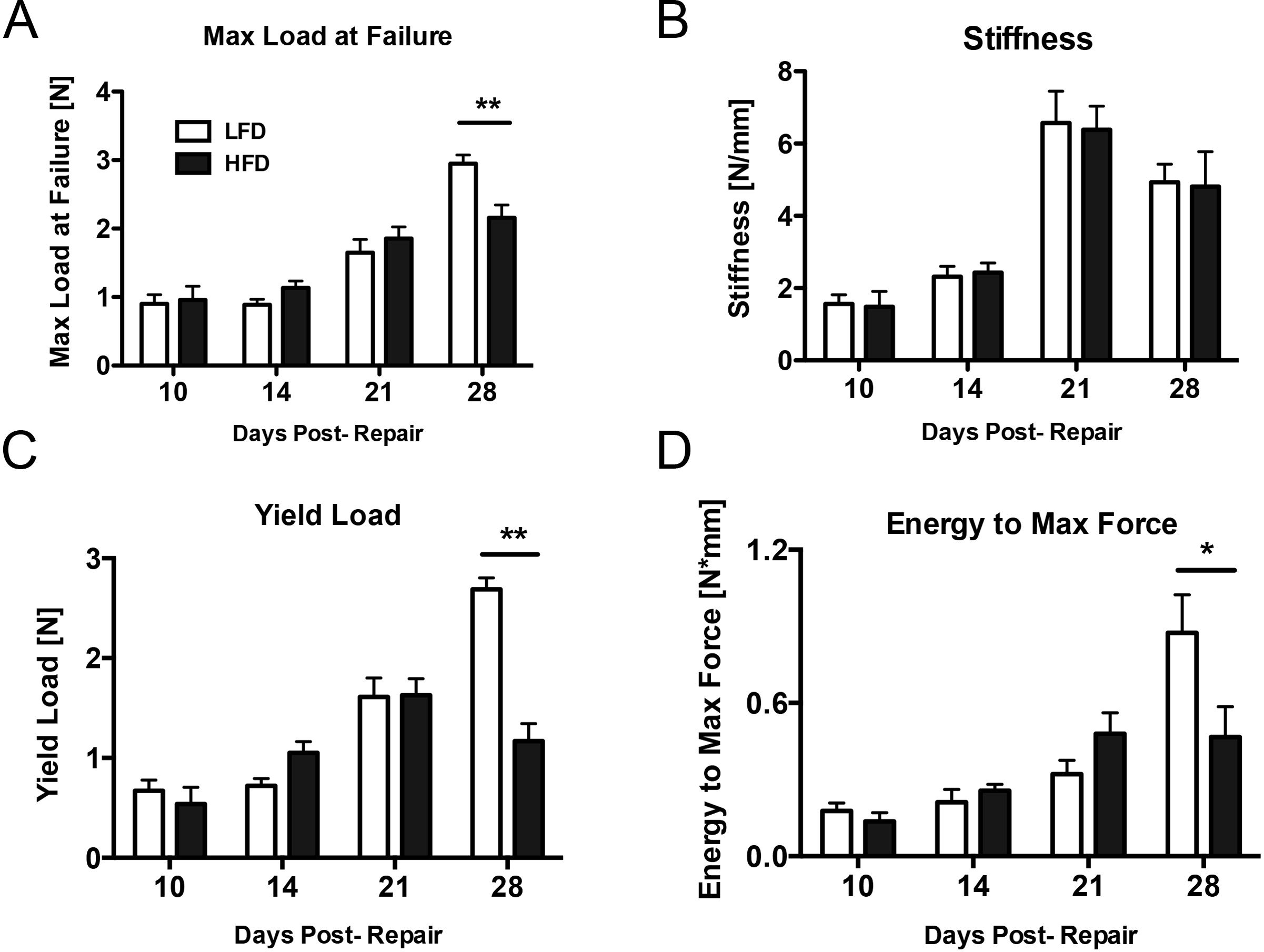
HFD impairs biomechanical properties of the tendon repair. Repaired tendons underwent tensile testing to failure at different time points after surgery to determine the effects of T2DM on mechanical properties of the tissue. A – The HFD group (black bars) had significantly lower maximum load at failure 28 days post-repair relative to the LFD group (white bars). B – Stiffness was not significantly different between HFD (black bars) and LFD (white bars) at any time after surgery. C – The yield load was significantly lower 28 days post-repair in the HFD group (black bars). D – Energy to maximum force was significantly lower 28 days post-repair in the HFD group (black bars) relative to the LFD group. (*) Indicates p<0.05, (**) indicates p<0.01.

### Adhesion remodeling is delayed in the HFD group

At 7 and 10 days both LFD and HFD repairs demonstrated a robust granulation response bridging the injury site (outlined in green) (Figure 4); this granulation response was especially robust at day 10 in the HFD group. Adhesions formed between the tendon (outlined in yellow) and soft-tissue at day 14 days in both groups, which were observed as a tight junction between tissue types (black arrowhead). However, LFD repairs demonstrated some space between the tendon and soft-tissue below the repair at 14 days (black arrow), suggesting fewer adhesions in this area. Substantial adhesions were present around the repair in the HFD group at 21 days (black arrowhead), while some separation between tissues was present at 21 days (black arrows) in both HFD and LFD repairs. By day 28, there was evident spacing both above and below the repair in LFD repairs (black arrows), while adhesions were still present in the HFD repairs (black arrowhead).

**Figure 4.**
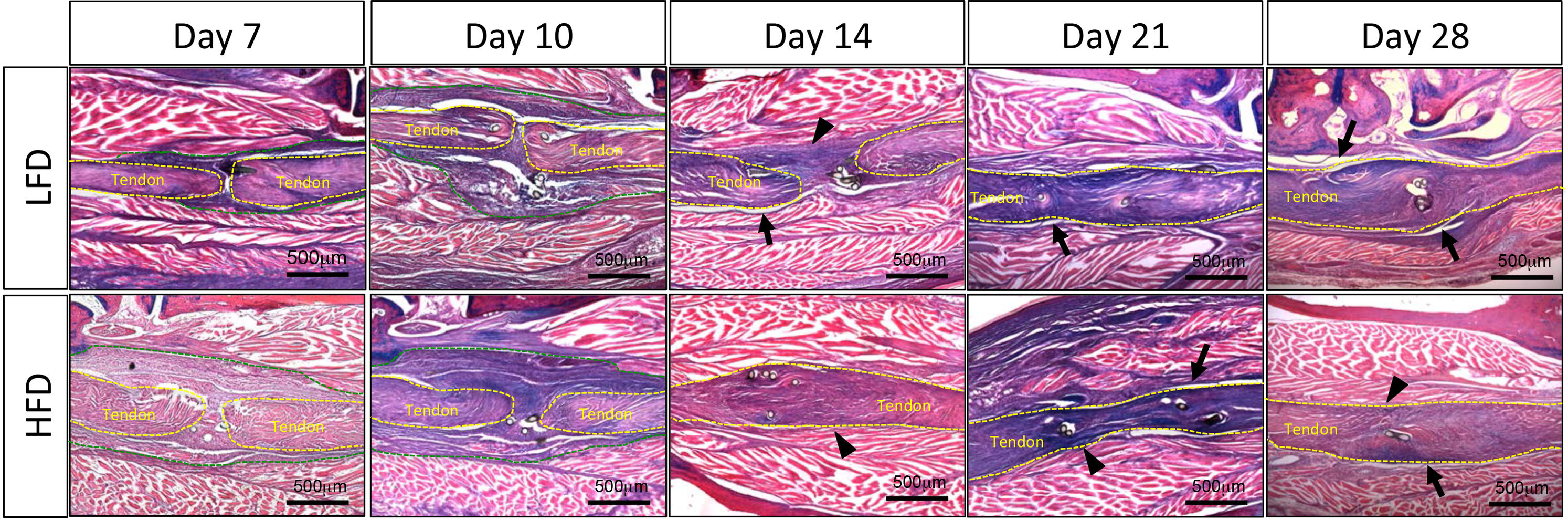
Adhesion remodeling is delayed in the HFD group. Histology was performed to visualize changes to the repair site over time in HFD and control groups. Both groups demonstrated a robust granulation response early after surgery, with adhesions appreciable 14 days post-repair (black arrowhead). A clear boundary was visualized inferior to the tendon in control repair after 14 days (black arrow). Adhesions persisted in the HFD group at 21 days and 28 days post-repair (black arrowheads), while boundaries superior and inferior to the tendon were well defined in the control group (black arrows). (Alcian blue/hematoxylin and Orange G stain; scale bars represent 500 microns; native tendon tissue outlined in yellow; granulation/scar tissue outlined in green.

### Obesity/T2DM Increases Collagen and Mmp gene expression during healing

The proliferative phase of tendon healing is characterized by increased type III collagen production (*Col3a1)*, which is eventually remodeled to the more organized type I collagen that comprises the majority of homeostatic tendon. While LFD repairs demonstrated a predictable peak and decline in *Col3a1* expression consistent with the timing of the proliferative phase of healing, the pattern of *Col3a1* expression in HFD repairs was both increased and prolonged, with significant increases in expression relative to LFD repairs at 10, 14 and 21 days (Day 14- LFD: 0.26 ± 0.04, HFD: 3.03 ± 0.95, p<0.0001)(Figure 5A). Only by 28 days did expression of *Col3a1* return to levels comparable to LFD repairs (LFD: 0.52 ± 0.01, HFD: 1.34 ± 0.07, p>0.05). A significant increase in *Col1a1* expression was observed in LFD repairs, relative to HFD at day 3, however between 10-21 days significant increases in *Col1a1* expression were observed in HFD repairs, relative to LFD (Day 21- LFD: 0.49 ± 0.03, HFD: 3.25 ± 0.30, p<0.0001)(Figure 5B).

**Figure 5.**
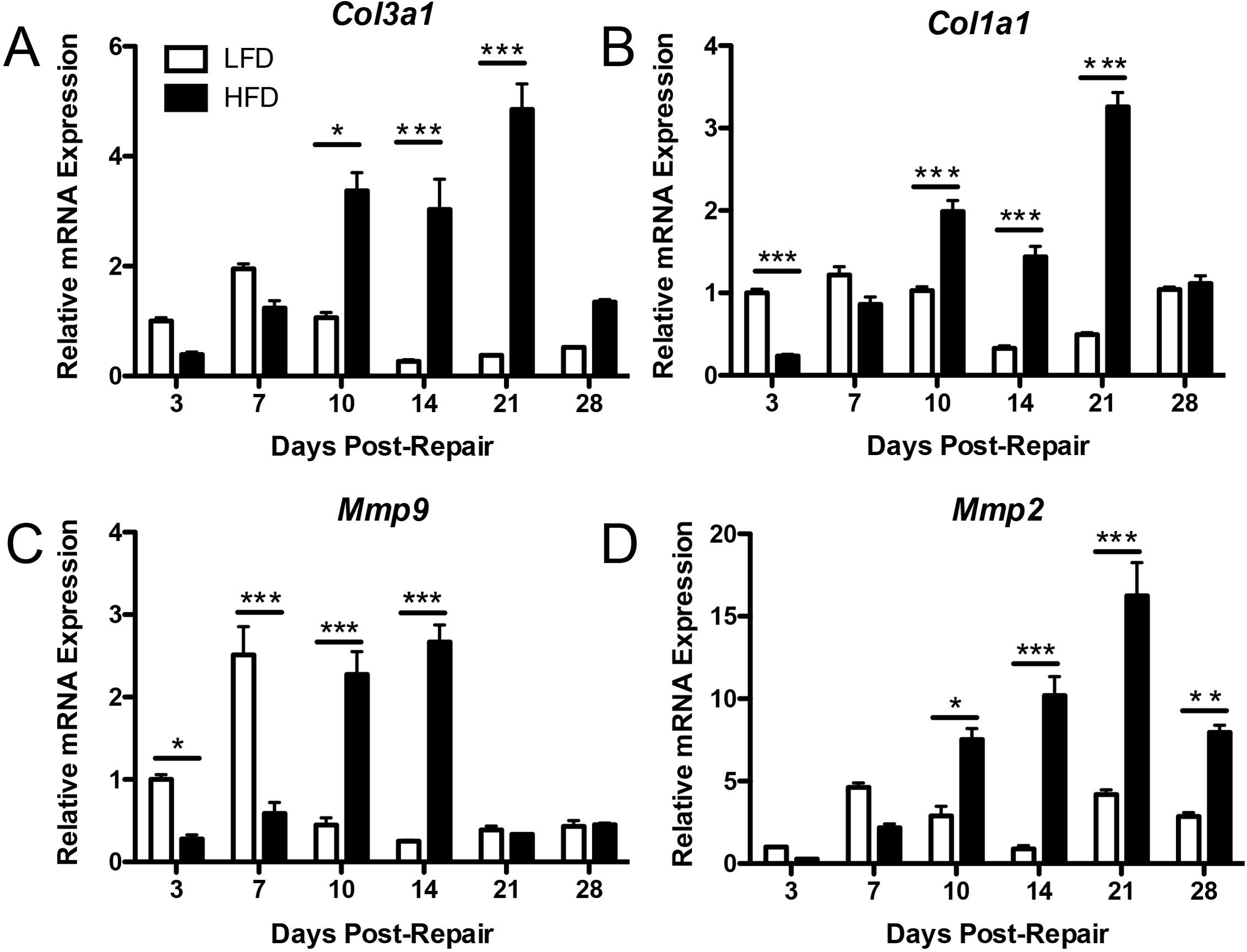
Relative expression of Collagen and Matrix Metalloproteinase genes are altered in HFD. Relative mRNA expression of (A) type III (*Col3a1*) and (B) type I collagen (*Col1a1*), (C) *Mmp9,* and (D) *Mmp2* were determined by RT-PCR between 3 and 28 days post-repair. Data were normalized to the internal control β*-actin*. Fold changes are reported relative to LFD day 3 expression. White bars = LFD; black bars = HFD, (*) indicates p<0.05, (**) indicates p<0.001, (***) indicates p<0.0001 between HFD and LFD.

*Mmp9* is associated with increased adhesion formation (17, 20, 21), while *Mmp2* expression is up-regulated during the remodeling phase of tendon repair (20). Peak *Mmp9* expression occurred day 7 in LFD repairs, while the HFD group demonstrated a delayed and significantly elevated expression pattern at 10 and 14 days (D10-LFD: 0.44 ± 0.14; HFD: 2.27 ± 0.47, p<0.0001) (D14- LFD: 0.25 ± 0.01; HFD: 2.66 ± 0.35, p<0.0001) (Figure 5C). Expression of *Mmp2* rose steadily from 3 to 21 days in the HFD group, with levels significantly higher than LFD between 10 and 28 days (Day 21- LFD: 4.18 ± 0.5, HFD: 16.23 ± 3.49, p<0.0001)(Figure 5D).

### The macrophage response to flexor tendon repair is altered in Obese/T2DM mice

Relative expression of *F4/80*, a pan-macrophage marker, was significantly elevated in the HFD group at 3, 7, 14 and 21 days relative to LFD (Day 7- LFD: 1.27 ± 0.31, HFD: 4.85 ± 0.24, p<0.001)(Figure 6A), suggesting an overall elevation in macrophage response to tendon injury in T2DM.

**Figure 6.**
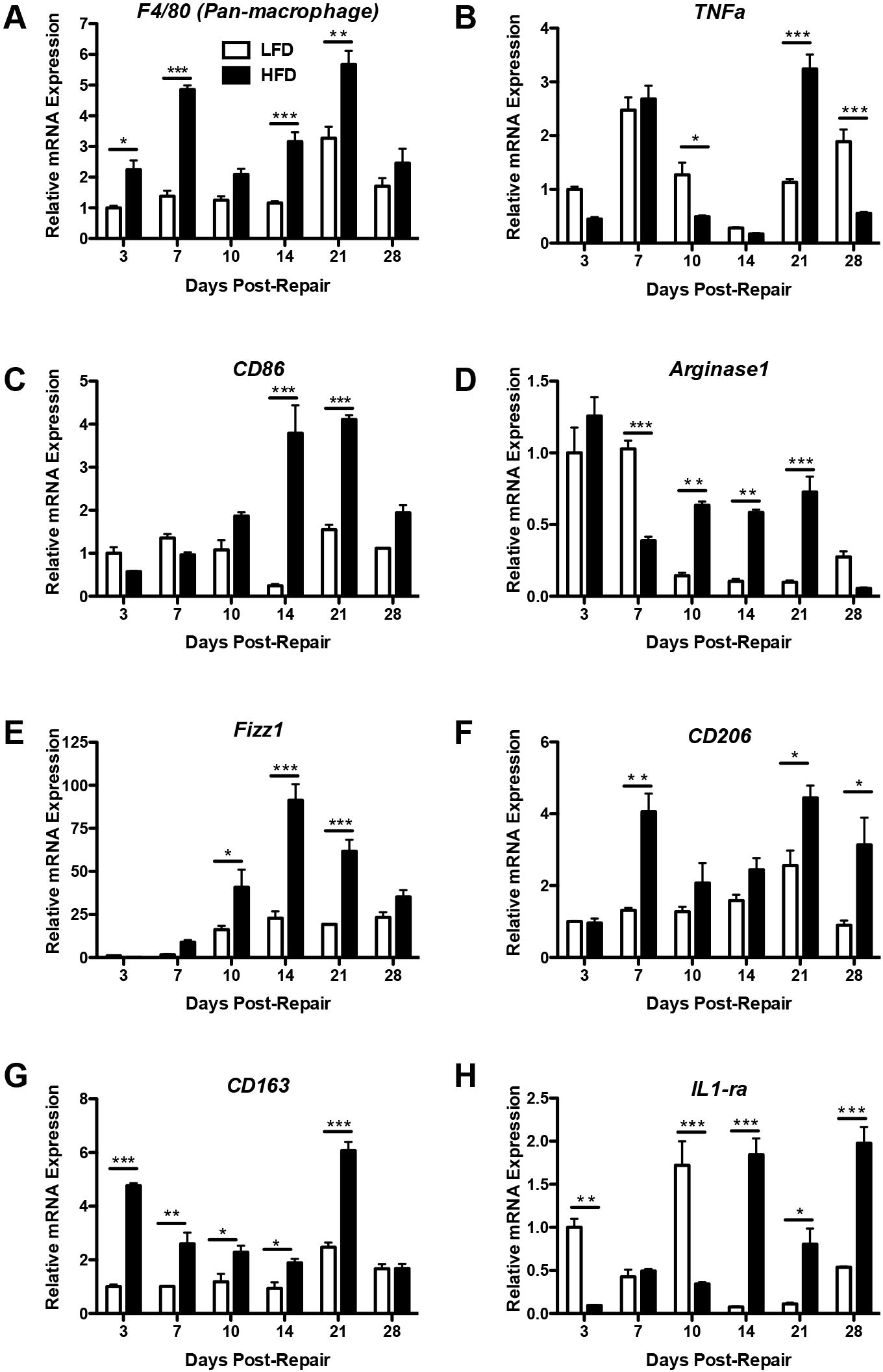
HFD alters macrophage polarization during tendon repair. Relative mRNA expression of (A) *F4/80;* M1 genes (B) *TNFa*, and (C) *CD86*; M2 genes (D) *Arginase1*, (E), *Fizz1* (F) *CD206*, (G) *CD163*, and (H) *IL-1ra* was determined by RT-PCR between 3 and 28 days post-repair. Data were normalized to the internal control β*-actin*. Fold changes are reported relative to LFD day 3 expression. White bars = LFD; black bars = HFD, (*) indicates p<0.05, (**) indicates p<0.001, (***) indicates p<0.0001 between HFD and LFD.

Macrophages are broadly categorized into M1 and M2 types based on their pro-inflammatory and anti-inflammatory properties, respectively (17). Peak expression of *TNF*α, a stimulator of the pro-inflammatory M1 phenotype (22), occurred on day 7 in LFD (2.47 ± 0.40), however, peak expression was delayed in HFD repairs until day 21 (3.24 ± 0.45) (Figure 6B). *CD86*, an M1 marker (13), was significantly increased in the HFD group at 14, and 21 days relative to LFD (Day 21- LFD: 1.54 ± 0.19; HFD: 4.10 ± 0.17, p<0.0001) (Figure 6C).

M2 macrophages are associated with tissue repair and matrix deposition. Peak expression of the M2 marker *Arginase1* occurred at day 3 in both HFD and LFD. In LFD repairs, *Arginase1* expression remained elevated at day 7 (p<0.001 vs. HFD), but decreased after this time, resulting in significant increases in HFD repairs, relative to LFD, at 10, 14, and 21 days (Day 21-LFD: 0.09 ± 0.02, HFD: 0.72 ± 0.18, p<0.0001)(Figure 6D). *Fizz1* expression was significantly elevated in the HFD between 10-21 days, relative to LFD (Day 21- LFD: 19.17 ± 0.2, HFD: 61.67 ± 11.66, p<0.0001)(Figure 6E). *CD206* was expressed by resident tissue M2 macrophages (23). Peak *CD206* expression occurred at day 21 in both LFD and HFD repairs, however, a significant increase in *CD206* expression was observed in HFD repairs at this time (Day 21- LFD: 2.55 ± 0.72, HFD: 4.44 ± 0.6, p<0.05. Additionally, *CD206* levels remained elevated in HFD repairs at day 28, relative to LFD (p<0.05)(Figure 6F). *CD163* was significantly increased in the HFD group between 3-21 days (Day 21- LFD: 2.47 ± 0.29, HFD: 6.07 ± 0.56, p<0.0001)(Figure 6G). *IL-1ra* was significantly increased in HFD at 10, 14, 21, and 28 days relative to LFD (Day 28- LFD: 0.53 ± 0.01, HFD: 1.97 ± 0.32, p<0.0001) (Figure 6H).

To more specifically trace macrophage content in obese/ diabetic mice relative to lean controls, MaFIA mice were placed on either HFD or LFD. Obesity and impaired glucose tolerance were confirmed by a significant increase in body weight in HFD MaFIA, relative to LFD MaFIA (Supplemental Figure 1A), and significantly increased blood glucose levels during GTT (Supplemental Figure 1B).

To assess systemic macrophage levels, peripheral blood was isolated at immediately prior to surgery (Day 0) or on days 7, 14 and 21 post-surgery. Significant increases in the proportion of GFP^+^ macrophages were observed in peripheral blood mononuclear cells (PBMCs) of HFD on days 0, 7, and 14 post-surgery (+18.4% on day 14, p=0.02). By day 21 no difference in PBMC GFP^+^ macrophage content was observed (Supplemental Figure 2A). A significant increase in bone marrow GFP^+^ macrophages was observed on day 21 post-surgery in HFD MaFIA mice, relative to LFD MaFIA (+19.9%, p=0.0015) (Supplemental Figure 2B).

Localization of macrophages at the flexor tendon repair site was identified by GFP and F4/80 expression. A slightly increased area of positive staining was observed, for both GFP and F4/80, in HFD repairs relative to LFD at day 21 post-surgery (Figure 7A, Supplemental Figure 2C). Consistent with qPCR data, an observable increase in both TNFα and IL-1ra positive cells were observed in HFD repairs, relative LFD (Figure 7B & C).

**Figure 7.**
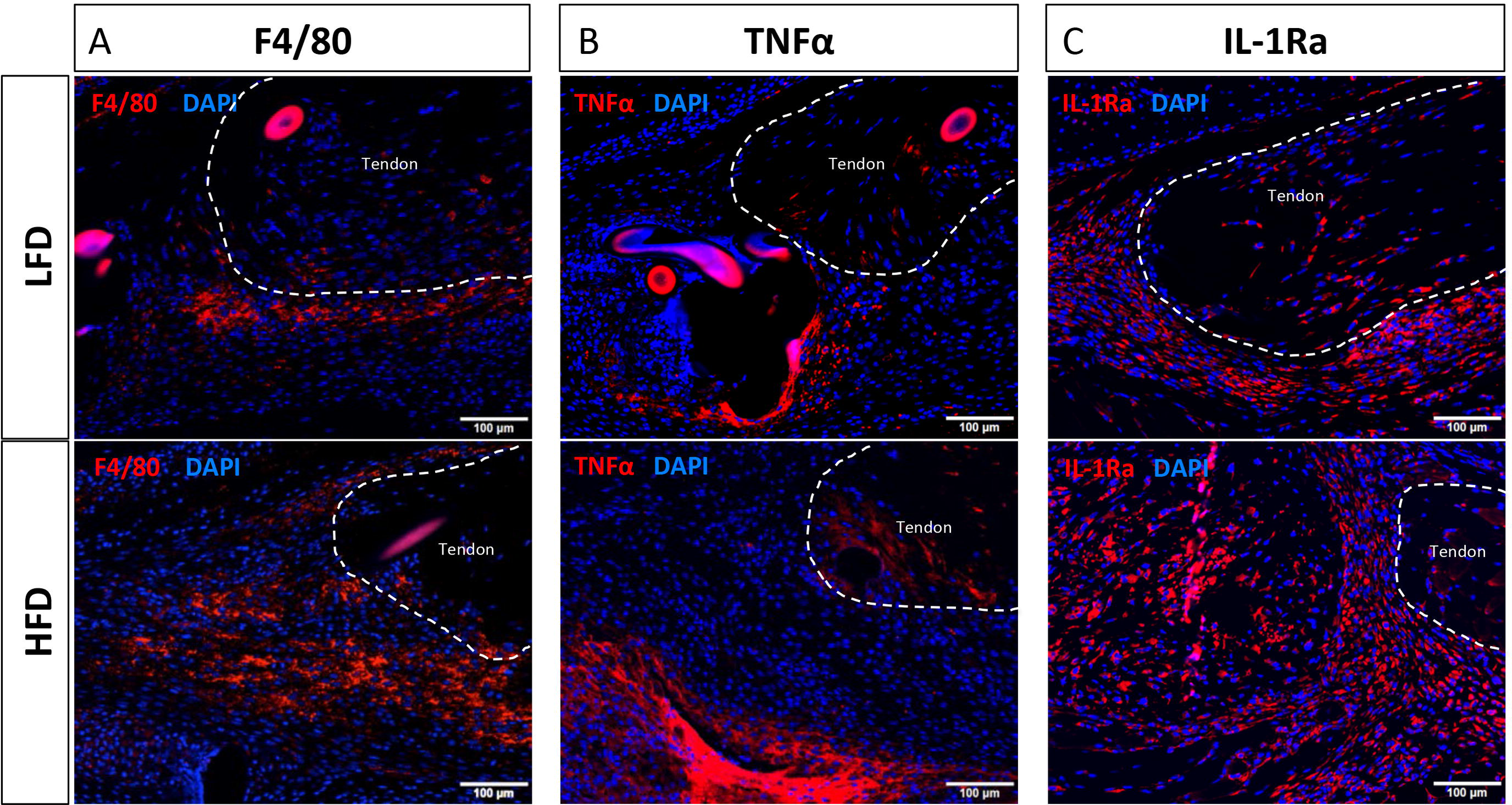
HFD tendons heal with increased TNF α and IL-1ra expression. Immunofluorescent staining for the pan-macrophage marker F4/80, the M1 macrophage marker TNFα, and the M2 macrophage marker IL-1ra at 21 days post-surgery in LFD and HFD repairs. Positive staining is indicated by red fluorescence, while the nuclear counterstain DAPI is blue. Native tendon is outlined in white. Scale bars represent 100μm.

## Discussion

Type II diabetes alters baseline tendon function and impairs the normal tendon healing process. Clinically, diabetic tendons have decreased ROM and are more likely to re-tear or rupture following repair (24), suggesting that T2DM alters tissue organization and quality. Animal models have demonstrated decreased mechanical properties (7), and decreased collagen fiber organization (8) in diabetic tendons relative to non-diabetic controls. However, little is understood about the mechanisms that govern impaired healing in type II diabetics, as such there is currently no therapeutic approach to improve outcomes in these patients. In the present study we have demonstrated that repaired FTs of obese/diabetic mice heal with prolonged scar formation, compromised mechanical properties, and increased deposition of extracellular matrix. Furthermore, elevated expression of markers associated with both the pro-inflammatory M1 macrophage phenotype, and the matrix deposition/ remodeling M2 macrophage are observed in obese/ diabetic repairs suggesting that an altered macrophage response may be associated with impairments in tendon healing that occur in type II diabetics.

Despite the clinical burden of impaired healing in diabetic patients, few studies have examined this scenario in animal models. Using non-obese type II diabetic Goto-Kakizaki (GK) rats, Ahmed *et al*., demonstrated a differential inflammatory profile in GK Achilles tendon repairs, relative to non-diabetic Wistar repairs, suggesting impaired regulation of inflammation in diabetic tendon repairs (9). Additionally, Bedi *et al*., demonstrated impaired tendon-bone healing in type I diabetic rats (25). However, Connizzo *et al*., established that different tendons exhibit unique responses to T2DM (4), while we recently demonstrated these same tendon-specific effects of obesity/ diabetes in additional tendons (5), suggesting that T2DM may differentially affect healing in other tendons. As such, characterization of the effects of T2DM specifically on the flexor tendon is important. Recently, David *et al*., established that the strength of murine obese/T2DM FT is decreased, possibly due to a decrease in cellularity relative to lean controls using a mid-substance biopsy punch model of healing (6). However, the effects of obesity/T2DM on gliding function were not assessed in that study. Assessing alterations in gliding function due to obesity/ diabetes is quite important given that gliding function is progressively diminished over the course of disease duration, however, no effects on gliding function are observed 12 weeks after diet initiation (5) suggesting that obesity/ T2DM does not perturb tendon homeostasis at the functional or mechanical level over the time period we have used in this study. The impairments in gliding function observed in obese/ diabetic mice suggest prolonged presence of scar tissue and adhesions in obese/T2DM repairs. Given that unsatisfactory digit ROM is one of the greatest complications of FT repairs (8), understanding the effects of obesity and T2DM on this process are particularly important.

Consistent with prolonged scar formation, obese/T2DM FT repairs also display significant increases in type III and type I collagen during the course of healing. Type III collagen is the predominant component of scar tissue and is associated with impaired mechanics. Excess matrix deposition can occur in several diabetic pathologies, including diabetic retinopathy and diabetic cardiomyopathy (11). While the precise mechanisms differ between tissues, chronic hyperglycemia and the subsequent tissue response are believed to underlie these fibrotic changes, with aberrant or excess deposition of type III collagen occurring in many diabetic pathologies (26).

While excess ECM deposition may result in increased scar formation in diabetic tendons, changes in matrix remodeling must also be considered. The effects of obesity/T2DM on Mmp expression and activity are not clear, and are likely tissue dependent. Muscles from diabetic mice have impaired Mmp activity (26), while Mmps are up-regulated in diabetic kidneys (27). Moreover, plasma Mmp levels were decreased in obese humans (28), while increased Mmp2 expression has been observed in adipose tissue (29). Here we show that both *Mmp9*, involved in degradation of damaged collagen, and *Mmp2*, associated with remodeling of disorganized ECM and scar tissue, are elevated in obese/T2DM repairs. Taken together, the collagen and Mmp expression data suggest that obese/diabetic tendons may heal with exuberant matrix deposition and remodeling. However, future studies should also evaluate changes in Mmp activity and assess healing and gliding function in diabetic repairs at later time-points to determine if the increase in Mmp expression may eventually restore gliding function in obese/diabetic repairs.

As macrophages play instrumental roles as both direct contributors and orchestrators of the inflammatory and reparative processes, defining their function in tendon repair is crucial to a better understanding of this process. At least two studies have demonstrated temporal changes in macrophage populations during Achilles tendon healing (30), further supporting a role for macrophages in both early inflammatory events and later tissue repair/ remodeling. Given that T2DM results in elevated systemic inflammation, we hypothesized that healing in T2DM tendons would result in an altered macrophage response that may underlie the changes associated with FT repair in T2DM. Expression levels of *TNF*α, a stimulator of the anti-inflammatory M1 phenotype, and *CD86*, a cell surface receptor expressed by M1 cells, are increased in LFD repairs during early healing. These findings are consistent with previous studies demonstrating a predominance of M1 macrophages during the inflammatory phase (31, 32). In contrast, peak expression of both *TNF*α and *CD86* are delayed until 21 days in obese/T2DM, suggesting either a delay in or an extension of the M1 inflammatory phenotype phase in diabetic tendon healing.

Alternatively-activated M2 macrophages are associated with tissue repair and remodeling (31-33). Recently, Shen *et al*. have identified enhanced M2 macrophage activation (via adipose stem cell sheets) as a promising approach to improve FT healing (13). However, excessive and altered M2 activity can directly impair healing and lead to fibrosis (33). Tendon repairs in both HFD and LFD show increased expression of M2 markers during the later stages of healing, however obese/T2DM repairs had increased and sustained elevation of M2-associated genes, relative to LFD. *CD206,* expressed by resident tissue macrophages (34), was significantly increased at 21 and 28 days in obese/T2DM repairs, relative to LFD. Importantly, *CD206* can promote tissue fibrosis through fibroblast proliferation and up-regulation of TGF-β (24). *Fizz1* is a secreted protein (35) that can stimulate type 1 collagen expression (36) and is dramatically increased in obese/T2DM repairs between 10-21 days, concomitant with elevated *Col3a1* and *Col1a1* expression in obese/T2DM repairs. These data identify exuberant M2 macrophage function as a potential mechanism of altered healing in T2DM, and suggest that a greater understanding and characterization of these cells will be crucial to improving outcomes in this population.

While these data recapitulate many clinical phenotypes associated with healing in diabetic patients, there are several limitations in this study that must be considered. First, we have used a murine model of FT healing that is not a true representation of the anatomy in Zone II of the human hand. Our injury is induced in the mid-portion of the paw, in an area where the synovial sheath is not present; as such, our model is not able to account for the contribution of synovial sheath derived cells to healing and scar formation. Furthermore, release of the tendon at the myotendinous junction (MTJ) decreases tension on the repair site, a scenario that does not occur clinically. Thus, decreased tension on the repair site is likely to slightly alter the healing process. However, this release was uniformly applied to all mice in this study, and we observe some restoration of the MTJ by 7-10 days post-surgery, suggesting that the repaired tendon experiences some tension for much of the healing process. Additionally, we have used only male mice, as male C57Bl/6 mice are much more susceptible to T2DM induced by a HFD than female mice (37, 38). Given that females placed on a HFD typically develop obesity without becoming diabetic (39), future studies comparing healing in male and female HFD and LFD mice will be critical to dissect the relative importance of obesity, hyperglycemia, and Type II Diabetes during tendon healing, something that we are unable to do in this study. Interestingly, Bedi *et al*., have demonstrated that streptozotocin-induced type I diabetes impairs tendon-to-bone healing, including decreased collagen organization, suggesting hyperglycemia as the predominant factor in impaired tendon healing in diabetics (type I and type II). Finally, given the progressive declines in flexor tendon gliding function and ECM organization that are observed long-term in HFD mice (5), it would be interesting to determine if healing is further impaired in mice with long-term type II diabetes.

In the present study we have demonstrated that a murine model of diet induced obesity and T2DM results in excess and prolonged scar tissue formation, decreased strength, and an altered macrophage response. Taken together, these data suggest that altered macrophage polarization, specifically over abundant M2 activity may lead to an excess matrix deposition phenotype in T2DM. This altered macrophage polarization may drive prolonged scar formation and compromised biomechanical properties. As such, these studies identify pro-fibrotic macrophages as a potential target to improve healing in T2DM patients.

## Acknowledgements

Author contributions: Study conception and design: AEL; Acquisition of data: JEA, MBG, CAO, FB; Analysis and interpretation of data: JEA, MBG, AEL; Drafting of manuscript: JEA, MBG, AEL; Revision and approval of manuscript: JEA, MBG, CAO, FB, AEL.

We would like to thank the Histology, Biochemistry and Molecular Imaging (HBMI) core for technical assistance with histology, and the Biomechanics and Multimodal Tissue Imaging (BMTI) Core for technical assistance with the biomechanical testing.

## Supplemental Figure Legends

**Supplemental Figure 1- HFD results in obesity/ T2DM in MaFIA mice.** A – Body weights from MaFIA mice in both HFD (black bars) and LFD groups (white bars) were compared at the time of surgery, with HFD MaFIA mice weighing significantly more than LFD. C – Mice from the HFD MaFIA group (black line) demonstrated an impaired ability to respond to glucose loading relative to LFD MaFIA (red line), as shown by significantly elevated blood glucose levels at 15, 30, 60, and 120 minutes after delivering a glucose bolus. (*) Indicates p<0.05.

**Supplemental Figure 2- Obesity/ T2DM increases systemic but not local GFP^+^ macrophages in MaFIA mice following tendon repair.** To assess changes in macrophage content in (A) peripheral blood mononuclear cells (PBMCs), (B) bone marrow, and (C) the healing flexor tendon, MaFIA mice, which express GFP in macrophages were used. Macrophage content was assessed via Flow cytometric analysis for GFP^+^ cells, in (A) Peripheral blood mononuclear cells (PBMCs) at days 0, 7, 14, and 21 days post-surgery, and (B) whole bone marrow at 21 days post-surgery. (*) Indicates p<0.05, (***) indicates p<0.0001 between HFD and LFD. No observable change in GFP expression was seen between LFD and HFD repairs, via GFP immunohistochemistry at day 21 post-surgery. Scale bar represents 100μm.

